# PYL8 ABA receptors of *Phoenix dactylifera* play a crucial role in response to abiotic stress and are stabilized by ABA (118)

**DOI:** 10.1101/2020.09.29.318170

**Authors:** Irene Garcia-Maquilon, Alberto Coego, Jorge Lozano-Juste, Maxim Messerer, Carlos de Ollas, Jose Julian, Rafael Ruiz-Partida, Gaston Pizzio, Borja Belda-Palazón, Aurelio Gomez-Cadenas, Klaus F.X. Mayer, Dietmar Geiger, Saleh A. Alquraishi, Abdulwahed F. Alrefaei, Peter Ache, Rainer Hedrich, Pedro L. Rodriguez

## Abstract

The identification of those prevailing ABA receptors and molecular mechanisms that trigger drought adaptation in crops well adapted to harsh conditions such as date palm (*Phoenix dactylifera*, Pd) sheds light on plant-environment interactions. We reveal that PdPYL8-like receptors are predominantly expressed under abiotic stress, being Pd27 the most expressed receptor in date palm. Therefore, subfamily I PdPYL8-like receptors have been selected for ABA signaling during abiotic stress response in this crop. Biochemical characterization of PdPYL8-like and PdPYL1-like receptors revealed receptor- and ABA-dependent inhibition of PP2Cs, which triggers activation of the *pRD29B-LUC* reporter in response to ABA. PdPYLs efficiently abolish PP2C-mediated repression of ABA signaling, but loss of the Trp lock in the seed-specific AHG1-like phosphatase PdPP2C79 markedly impairs its inhibition by ABA receptors. Characterization of *Arabidopsis* transgenic plants that express PdPYLs shows enhanced ABA signaling in seed, root and guard cells. Specifically, Pd27 overexpressing (OE) plants showed lower ABA content and were more efficient than wild type to lower transpiration at negative soil water potential, leading to enhanced drought tolerance. Finally, PdPYL8-like receptors accumulate after ABA treatment, which suggests that ABA-induced stabilization of these receptors operates in date palm for efficient boosting of ABA signaling in response to abiotic stress.

**Highlight:** Date palm response to abiotic stress is triggered through PYL8-like ABA receptors that are stabilized by the hormone, which boosts plant adaptation to drought mediated by ABA.

## Introduction

Among monocotyledons, the palm family plants (Arecaceae or Palmae) is second in economic importance after the grass family (Poaceae), and among the palm crops, the top three palm species are African oil palm (*Elaeis guineensis*), date palm (*Phoenix dactylifera*) and coconut (*Cocos nucifera*). Date palm (*Phoenix dactylifera* L., 2n=36) is thought to be native to the Arabian Peninsula from where was introduced to northern India, North Africa and southern Spain (Al-Mssallem et al., 2013). Date palm used to be the main nutrient for the population of Arabian Peninsula and currently is a major food crop in the Middle East, northern Africa and other places with suitable climate (Yin et al., 2012). Additionally, it is a sociocultural symbol in many countries with long agriculture history, being one of oldest domesticated trees (Al-Mssallem et al., 2013).

Date palm is a plant species adapted for growth under dry conditions and recent studies indicate that heat and drought stresses only slightly affect photosynthesis (Arab et al., 2016). Fast guard cell abscisic acid (ABA) signaling is present in date palm to limit transpirational water loss and the *Phoenix dactylifera* (Pd) SLAC1-type anion channel is regulated by the ABA kinase PdOST1 (Müller et al., 2017). In the presence of nitrate at the extracellular face of the anion channel, ABA enhanced and accelerated stomatal closure (Müller et al., 2017). Additionally, the date palm genome displays an expanded late embryogenesis abundant (LEA) gene family, whose ABA-induced expression is important for resistance to abiotic stress, particularly LEA2 genes that are ubiquitously expressed (Al-Mssallem et al., 2013). Given that ABA plays a key role for plant adaptation to drought and date palm is well adapted to survive and produce fruits under harsh environmental conditions, the characterization of core ABA signaling components in Pd is of great interest. ABA signaling is initiated by ABA perception through PYRABACTIN RESISTANCE1 (PYR1)/PYR1-LIKE (PYL)/REGULATORY COMPONENTS OF ABA RECEPTORS (RCAR) receptors (Park et al., 2009; Ma et al., 2009; Santiago et al., 2009; Rodriguez et al., 2019), which leads to interaction with and inactivation of clade A protein phosphatase type 2Cs (PP2Cs), such as ABA INSENSITIVE 1 (ABI1) and ABI2, HYPERSENSITIVE TO ABA (HAB1) and HAB2, and PROTEIN PHOSPHATASE 2CA/ABA-HYPERSENSITIVE GERMINATION 3 (PP2CA/AHG3), thereby relieving their inhibition on three ABA-activated SNF1-related protein kinases (SnRK2s) termed subclass III SnRK2s, i.e. SnRK2.2/SnRK2D, 2.3/I and 2.6/E/OST1 (Umezawa et al., 2009; Vlad et al., 2009).

The discovery of the 14-member gene family of PYR/PYL/RCAR ABA receptors in *Arabidopsis thaliana* (Park et al., 2009; Ma et al., 2009; Santiago et al., 2009) has enabled the identification of the multigene families of ABA receptors in different crops. Functional characterization of crop ABA receptors is required to determine the prevailing receptors for abiotic stress adaptation; however, this knowledge is limited (Gonzalez-Guzman et al., 2014; Miao et al., 2018). Crop PYR/PYL/RCARs belong to multi-gene families, which are composed of 15 members in tomato (Gonzalez-Guzman et al., 2014), 12 in rice (He et al., 2014), 13 in maize (He et al., 2018), 9 in wheat (Mega et al., 2019), 9 in barley (Seiler et al., 2014) and 11 in sweet orange (Arbona et al., 2017). The receptors encoded in these genomes are distributed in three distinct subfamilies based on amino acid sequence identity, which are reminiscent of the *Arabidopsis* PYR1/PYL1-like (clade III), PYL4-like (clade II) and PYL8-like subfamilies (clade I) (Rodriguez et al., 2019). However, not all families play similar roles for ABA signaling in different tissues and specific functions for some members of these subfamilies have emerged in the last years (Antoni et al., 2013; Zhao et al., 2016; Dittrich et al., 2019). In particular, PYL8 plays a non-redundant role for ABA signaling in root, and ABA diminishes ubiquitination of this unique receptor, which leads to its stabilization and accumulation in presence of ABA (Belda-Palazon et al., 2018). In contrast to *Arabidopsis*, functional knowledge in crops is scarce and prevents further biotechnological use, for example, to regulate transpiration or root architecture in order to better adapt plants to drought.

The genome of date palm (estimated genome length 670 Mb) has been sequenced using different genome assemblies (Al-Dous et al., 2011; Al-Mssallem et al., 2013; Hazzouri et al., 2015 and 2019), and different transcriptomic studies have been published (reviewed by Gros-Balthazard et al., 2018). This race for genomic data has enabled the identification and characterization of the sex determination locus (Torres et al., 2018), pathways responsible for fruit maturation (Al-Mssallem et al., 2013; Hazzouri et al., 2019) and elucidation of the domestication history and diversity (Hazzouri et al., 2015; Gros-Balthazard et al., 2018). RNA-seq expression data are available for the different fruit stages, response to different stress treatments and comparative transcriptome analysis of oil palm and date palm mesocarp (Bourgis et al., 2011; Yin et al., 2012; Yaish et al., 2017; Al-Harrasi et al., 2018; Xiao et al., 2019, Hazzouri et al., 2019). Genomic resources have been used along this work to identify the Pd ABA receptor family and pinpoint the major Pd ABA receptors expressed during ABA treatment, salt, heat, drought or combined heat-drought conditions (Müller et al., 2017; Safronov et al., 2017). As a result, we reveal that the PYL8-like ABA receptors of *Phoenix dactylifera* play a major role in response to abiotic stress and their accumulation is promoted by ABA through a posttranscriptional mechanism. Functional characterization of Phoenix ABA receptors indicates that inhibition of clade A PP2Cs is crucial for drought tolerance of date palm.

## Materials and methods

### Plant material and growth conditions

Two-year-old date palm seedlings were cultivated at 22/16°C, 50% RH, 12h light regime. For ABA treatment, single cut pinnae were treated for 2h with CO2-free air in darkness to fully open stomata, and then illuminated for 4h at a photon flux rate of 620 μmol/m^2^s. Then 25 μM ABA was fed via the petiole. After 2h, RNA was extracted from the pinnae. Control was treated equally with ABA-free solution. RNA was extracted from leaves and RNA-seq library generation and sequencing was described by Müller et al., (2017). For heat, drought and heat+drought treatments, plants were maintained under greenhouse conditions and transferred to growth chambers for stress treatments. In growth chambers, plants were first acclimatized for two weeks (16/8 h photoperiod and 20/15°C, 70% RH, at a photon flux rate of 200-300 μmol/m^2^s at leaf level). Experiments were carried out in two batches. In the first batch, plants were irrigated every second day (well-watered) and exposed to different growth temperatures: 20/15°C (70% RH) or 35/15°C (60-70% RH) for two weeks, followed by harvesting 6 h after the onset of light. In the second batch, watering was stopped after the acclimation period (drought conditions) and plants grown at 35°C were harvested after 4-5 days (drought + heat) whereas plants grown at 20°C were collected after 7-8 days (drought). For salt treatment, pots were flooded with salt solution for 1 h every second day starting from day 2. After two weeks, RNA was extracted from plants treated with salt-free solution, 200 mM salt solution or 600 mM salt solution. RNA-seq library generation and sequencing was performed as described by Safronov et al. (2017). Root and fruit expression data for date palm have been reported by Xiao et al., (2019) and Hazzouri et al. (2019), respectively. Briefly, germinated root tips (from 12-week-old seedlings) were sampled for RNA extraction. Total RNA was extracted from 75 mg of plant material. Fruits of the Khalas variety were sampled at 45, 75, 105, 120 and 135 days after pollination (DAP). *Arabidopsis thaliana* plants were grown as described by Planes et al. (2015).

### Transpiration assay

Transpiration was measured in 16 plants per genotype that were grown in individual jiffy peat plugs (Jiffy-7 peat pellets, Semillas Batlle S.A., Barcelona, Spain) under different soil water potential conditions according to de Ollas et al., (2019). Plants were kept under well-watered conditions for 4 weeks in a growth chamber (Equitec model EGCS 351 3S HR, Getafe, Spain) with a day/night temperature of 23/18 °C, an 8 h light photoperiod (100 μmol m^−2^ s^−1^), and a relative humidity of 60–65%. To set up the drought conditions plants were randomly divided in 4 groups of 4 plants and differentially watered to cover the soil water content range from well-watered conditions to plant wilting (de Ollas et al., 2019). To characterize transpiration, individual plugs were weighed every 24 hours to account for plants transpiration water loss and individually normalized to leaf area. Transpiration was measured two consecutive days before harvesting. Plant tissue of each individual plant was immediately frozen in liquid N2 and lyophilized to analyze ABA endogenous contend.

### ABA quantification

ABA analysis was performed according to Durgbanshi, A. et al., (2005) with slight modifications. Briefly, 0.1 g of dry plant material was extracted in 2 ml of distilled H_2_O after spiking with 25 μl of a 2 mg l^−1^ solution of d6-ABA as internal standard. After centrifugation (10,000g at 4 °C), supernatants were recovered, and the pH was adjusted to 3.0 with 30% acetic acid. The acidified water extract was partitioned twice against 3 ml of di-ethyl ether. The organic layer was recovered and evaporated under vacuum in a centrifuge concentrator (Speed Vac, Jouan, Saint Herblain Cedex, France). The dry residue was then re-suspended in a 9:1 H_2_O:MeOH solution by sonication. The resulting solution was filtered and directly injected into a UPLC system (Waters Acquity SDS, Waters Corp., Milford, MA, USA) interfaced to a TQD triple quadrupole (Micromass Ltd, Manchester, UK) mass spectrometer through an orthogonal Z-spray electrospray ion source. Separations were carried out on a Gravity C18 column (50□×□2.1 mm, 1.8 μm, Macherey–Nagel GmbH, Germany) using a linear gradient of MeOH and H2O supplemented with 0.1% acetic acid at a flow rate of 300 μl min^−1^. Transitions for ABA/d6-ABA (263□>□153/269□>□159) were monitored in negative ionization mode. Quantitation of plant hormones was achieved by external calibration with known amounts of pure standards using Masslynx v4.1 software.

### Generation of Arabidopsis transgenic plants

The indicated pAlligator2-*35S:Pd* and *pMDC43-35S:GFP-Pd* constructs were transferred to *Agrobacterium tumefaciens* C58C1 (pGV2260) (Deblaere *et al.*, 1985) by electroporation and used to transform Columbia wild type plants by the floral dip method (Clough and Bent, 1998). T1 transgenic seeds were selected based on seed GFP fluorescence (pAlligator2) or hygromicin resistance (pMDC43), and sowed in soil to obtain the T2 generation. Homozygous T3 progeny was used for further studies and expression of HA-tagged or GFP tagged protein was verified by immunoblot analysis using anti-HA-HRP or anti-GFP antibodies, respectively.

### Constructs

The PdPP2C55, PdPP2C79, Pd15, Pd27, Pd32, Pd44, Pd957 coding sequences were obtained as synthetic DNA fragments (Invitrogen), which were amplified using the indicated primers (Supplementary Table S1), verified by sequencing and cloned into pCR8/GW/TOPO. In order to obtain recombinant protein, each coding sequence was excised using an *NcoI-EcoRI* double digestion and cloned into pETM11.

### Chemicals

The ABA agonists used in this work were provided by Dr Sean R. Cutler (University of California, Riverside, USA). ABA was obtained from Biosynth (https://www.biosynth.com/).

### Transient protein expression in Nicotiana benthamiana

*Agrobacterium* infiltration of *N. benthamiana* leaves was performed basically as described by Saez et al. (2008). For transient expression of GFP-PdPP2C55, PdPP2C79 or Pd ABA receptors, the corresponding pMDC43 binary vector was introduced into *A. tumefaciens* C58C1 (pGV2260) by electroporation and transformed cells were selected in LB plates supplemented with kanamycin (50 ug/ml). Then, they were grown in liquid LB medium to late exponential phase and cells were harvested by centrifugation and resuspended in 10 mM morpholinoethanesulphonic (MES) acid-KOH pH 5.6 containing 10 mM MgCl2 and 150 mM acetosyringone to an OD600 nm of 1. These cells were mixed with an equal volume of *Agrobacterium* C58C1 (pCH32 35S:p19) expressing the silencing suppressor p19 of tomato bushy stunt virus so that the final density of *Agrobacterium* solution was about 1 (final concentration OD600=0.5 each). Bacteria were incubated for 3 h at room temperature and then injected into young fully expanded leaves of 4-week-old *N. benthamiana* plants. Leaves were examined 48-72 h after infiltration using confocal laser scanning microscopy.

### Protoplast transfection

We analyzed ABA signaling in *Arabidopsis* protoplasts basically as described by Fujii et al. (2009). Briefly, protoplasts prepared from wild-type Col-0 plants were transfected with the reporter construct pRD29B:LUC, p35S:GUS for normalization, and effector plasmids encoding ABF2, ABF2+HAB1 or ABF2+HAB1+PdPYL proteins. LUC activity was measured in protein extracts prepared from protoplast suspensions 6 h after transfection, either in the absence or presence of 5 μM exogenous ABA that was added 3 h before measuring LUC activity. The activity of the pRD29B:LUC reporter was normalized with p35S:GUS as described by Fujii et al. (2009). To generate the effector plasmids, the coding sequence of ABF2, HAB1, Pd15, Pd27, Pd32, Pd44 or Pd957 was recombined by LR reaction from pCR8 entry vector to pXCS destination vector (Witte et al., 2004). The pRD29B:LUC and 35S:GUS pSK constructs were previously described (Christmann et al., 2005; Moes et al., 2008)

### Protein extraction, analysis and immunodetection

Protein extracts for immunodetection experiments were prepared from *Arabidopsis* transgenic lines expressing HA-tagged or GFP-tagged ABA receptors or GFP-tagged PP2C/ABA receptors transiently expressed in *Nicotiana benthamiana* as described by Fernandez et al., (2020).

### Confocal Laser Scanning Microscopy (CLSM)

Confocal imaging was performed using a Zeiss LSM 780 AxioObserver.Z1 laser scanning microscope with C-Apochromat 40x/1.20 W corrective water immersion objective. The GFP fluorophore was excited and fluorescence emission detected at the indicated wavelengths: 488 nm/500-530 nm, and chlorophyll at 561 nm/ 685-760 nm. Post-acquisition image processing was performed using ZEN (ZEISS Efficient Navigation) Lite 2012 imaging software and ImageJ (http://rsb.info.gov/ij/).

### Seed germination and seedling establishment assays

After surface sterilization of the seeds, stratification was conducted in the dark at 4°C for 3 d. Approximately 100 seeds of each genotype were sowed on MS plates supplemented with different ABA concentrations per experiment. To score seed germination, radical emergence was analyzed at 72 h after sowing. Seedling establishment was scored as the percentage of seeds that developed green expanded cotyledons and the first pair of true leaves at 5 or 7 d. Seedling establishment assays in the presence of ABA agonists were performed in 24-well plates, where approximately 25 seeds of the indicated genotype (three independent experiments) were sown on wells lacking or supplemented with the indicated concentration of ABA, QB or AMF4.

### Root growth assay

Seedlings were grown on vertically oriented Murashige and Skoog (MS) plates for 4-5 days. Afterwards, 20 plants were transferred to new MS plates lacking or supplemented with 10 μM ABA. The plates were scanned on a flatbed scanner after 10 days to produce image files suitable for quantitative analysis of root growth using the NIH Image software ImageJ

### Drought resistance experiments

Three-weeks-old plants (n=10, three independent experiments) were grown under greenhouse conditions (40-50% room humidity) and standard watering for 15 d. Watering was withheld for 20 d (Pd27 OE lines) or 8 d (PdPP2C79 OE lines), and then restored. Survival of the plants was scored 5 d later.

### Infrared thermography

Plants were grown in a controlled environment growth chamber at 22°C under a 12 h light, 12 h dark photoperiod at 100 μE m-2 sec-1 and 40-50% room humidity. Philips bulbs were used (TL-D Super 8036W, white light 840, 4000K light code). Infrared thermography images of rosette leaves were acquired from 6 week-old plants with a thermal camera FLIR E95 equipped with a 42° lens. Images were processed and quantified with the FLIR tools software. For quantification, the average temperature of 15 different sections corresponding to 4 leafs per plant were calculated. 5 plants per genotype were analyzed in each experiment. The mean temperature ± standard deviation of all the plants for each genotype was reported. Statistical comparisons among genotypes were performed by pairwise t-tests.

### PP2C inhibition assays

The expression in bacteria and purification of 6His-ΔNHAB1, PdPP2C55, PdPP2C79, Pd15, Pd27, Pd32, Pd44 and Pd957 was performed as described by Santiago et al. (2009). His-tagged proteins were purified using Ni-NTA affinity chromatography, eluted and analysed by SDS-PAGE, followed by Instant Blue staining. Phosphatase activity of ΔNHAB1 was measured using pNPP (15 mM) as substrate, 1 μM of the PP2C and 2 μM of the indicated receptors. Dephosphorylation of pNPP was monitored with a ViktorX5 reader at 405 nm (Antoni et al., 2012). Phosphatase activity of PdPP2C55 and PdPP2C79 was measured using the RRA(phosphoT)VA peptide as substrate, which has a Km of 0.5-1 μM for eukaryotic PP2Cs. Assays were performed in a 100 μl reaction volume containing 25 mM Tris–HCl pH 7.5, 10 mM MgCl2, 1 mM DTT, 25 μM peptide substrate and 1 μM of the PP2C and 2 μM of the indicated receptors. After incubation for 60 min at 30°C, the reaction was stopped by addition of 30 μl molybdate dye (Baykov et al., 1988) and the absorbance was read at 630 nm with a 96-well plate reader.

### Native gel electrophoresis

Native Red Electrophoresis was carried out as described previously (Ruiz-Partida et al., 2018). Protein samples, each containing 8.5 μg of purified protein in sample buffer: 50 mM Tris–HCl (pH 7.5), 100 mM NaCl, 5 mM MgCl2, 15 % glycerol and 0.02 % Red Ponceau 2S, with 50 mM DTT, were loaded onto 15 % polyacrylamide gels prepared in 375 mM Tris–HCl (pH 8.8), 10 % glycerol, 0.012 % of Ponceau Red S. A solution of Tris–Glycine (pH 8.8) with or without 0.012 % Ponceau Red S was used as the cathode and anode buffer, respectively. Electrophoresis was performed at a constant current of 25 mA for 150 min at 4 °C. Finally, proteins were detected in gel using standard Coomassie Blue staining.

## Results

### The date palm genome encodes 12 putative PYR/PYL/RCAR ABA receptors

We did a BLAST search in Kyoto Encyclopedia of Genes and Genomes (KEGG) database (https://www.genome.jp/kegg-bin/show_organism?org=pda) to identify PYR/PYL/RCAR Pd ABA receptors using as a query the *Arabidopsis* receptors PYL1, PYL4 and PYL8. As a result, we found a family composed of 12 Pd ABA receptors (Fig. 1A). With the exception of 91, they were distributed in three subfamilies, which match the corresponding groups from *Arabidopsis* receptors. This enables a tentative translation to Pd ABA receptors of those biochemical and physiological features that are already known in *Arabidopsis* ABA receptors (Rodriguez et al., 2019). In subfamily III we found three Pd receptors that show sequence similarity with dimeric AtPYL1, whereas in both subfamilies II and I we found four Pd receptors that are related to PYL4 and PYL8, respectively. Given that the *Arabidopsis* genome contains 14 receptors and date palm only 12, it is not possible to establish a direct correlation in the nomenclature with each *Arabidopsis* receptor. Therefore, we have preferred to maintain the KEGG-NCBI-Gene ID nomenclature to assign unambiguous names, using the last two or three digits of the full KEGG code to name each individual receptor (Fig. 1). Tentatively, some features of *Arabidopsis* PYL receptors can be assigned to Pd receptors based on sequence similarity and their location in different branches of the family. Amino acid sequence alignment of Pd ABA receptors (Fig. 1B) revealed the conserved residues of the gate (SGLPA) and latch (HRL) loops (defined by the b3-b4 and b5-b6 regions), which are a hallmark to define PYR/PYL/RCAR ABA receptors in different plant species (Rodriguez et al., 2019).

**Fig. 1.**
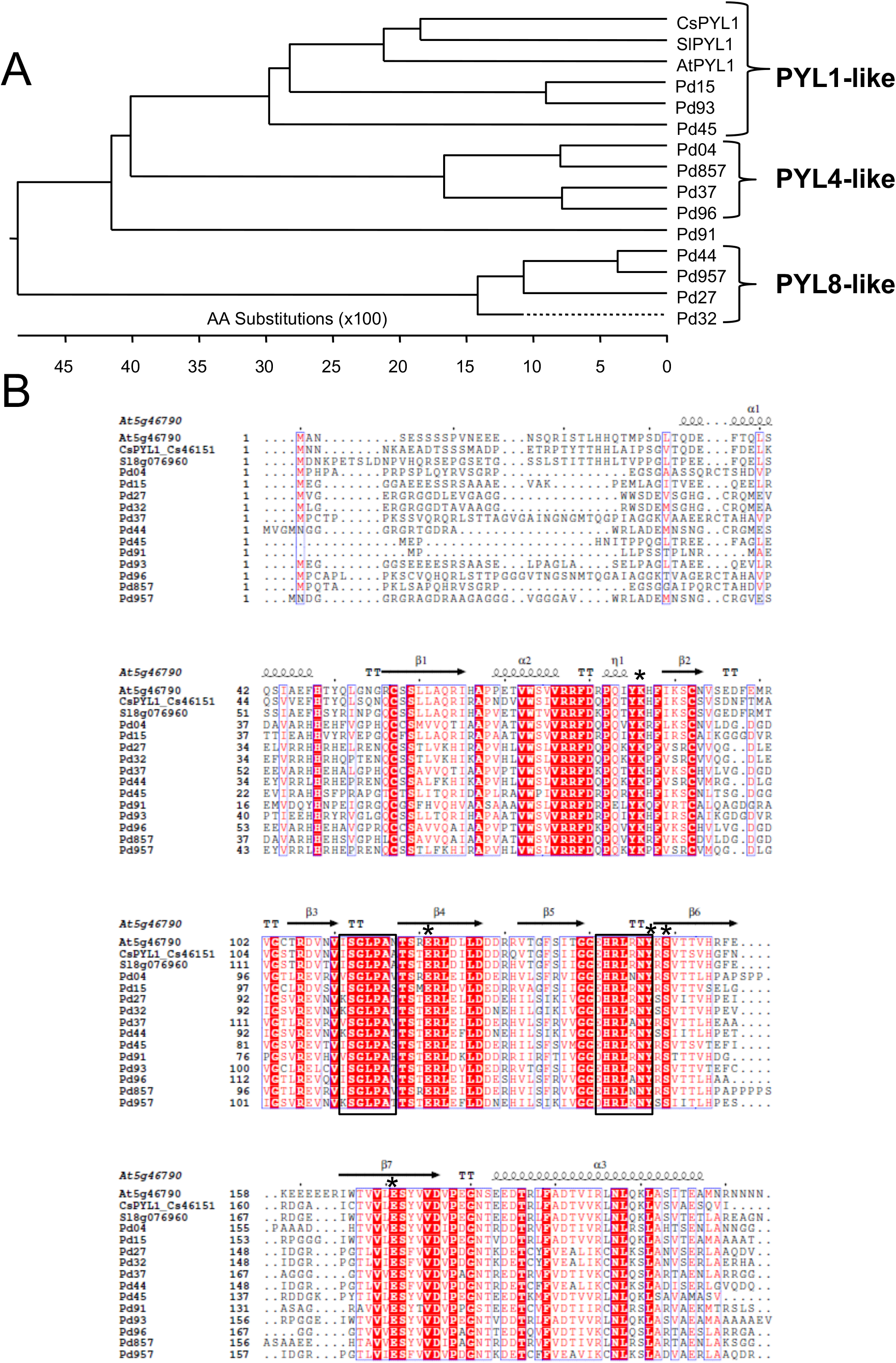
Cladogram and amino acid sequence alignment of date palm PYR/PYL/RCAR ABA receptors. (A) Cladogram of the multiple sequence alignment of date palm and *Arabidopsis* receptors, revealing three major subfamilies and the ungrouped Pd91. Sequences of *Citrus sinensis* (CsPYL1, Cs46151) and *Solanum lycopersicum* (SlPYL1, Sl8g076960) PYL1-like receptors were also included. The nomenclature is given from the last two or three digits of KEGG database entries for Pd genes: 1037118<u>15</u>, 1037046<u>93</u>, 1037095<u>45</u>, 1037144<u>04</u>, 1037088<u>57</u>, 1036964<u>37</u>, 1037174<u>96</u>, 1037068<u>91</u>, 1037071<u>44</u>, 103702<u>957</u>, 1036985<u>27</u> and 1037083<u>32</u>. The tree was constructed using GeneDoc and ClustalW software. (B) Sequence and secondary structure alignment of date palm ABA receptors and *Arabidopsis*, sweet orange and tomato PYL1 proteins. The predicted secondary structure of the date palm receptors is indicated, taking as a model the crystallographic structure of AtPYL1 (Protein DataBank Code 3JRS) and using the ESPRIPT program (http://espript.ibcp.fr/ESPript/ESPript/). Boxes indicate the position of the gate and latch loops. Black asterisks mark residues involved in interactions with ABA’s carboxylate group (K86, E121, Y147, S149 and E171 of AtPYL1), either direct contact for the Lys side chain or hydrogen bonds mediated by internal water molecules for the other residues.

### Transcription profiles of Pd ABA receptors reveal primacy of the PYL8-like family in abiotic stress response

Different RNA-Seq gene expression data for the Pd transcriptome have been published (Bourgis et al., 2011; Yin et al., 2012; Al-Mssallem et al., 2013; Yaish et al., 2017; Hazzouri et al., 2019; Xiao et al., 2019). We extracted transcription profiles of Pd ABA receptors from abiotic stress studies performed by Müller et al. (2017) and Safronov et al. (2017). Additionally, in this study we generated new RNA-seq data after mock- or salt treatment (200 and 600 mM) of two-year-old date palms. We screened the ArrayExpress archive (E-MTAB-5261) at EMBL-EBI (https://www.ebi.ac.uk/arrayexpress/submit/overview.html) in order to obtain gene expression profiles for the 12 Pd ABA receptors after different treatments (Fig. 2A). In *Arabidopsis*, at least some representative members of the PYR1/PYL1-like, PYL4-like and PYL8-like subfamilies are expressed to similar levels (Gonzalez-Guzman et al., 2012), but few analyses have been reported in crops (Gonzalez-Guzman et al., 2014). Interestingly, in a crop adapted to harsh conditions as date palm, we found that four PYL8-like and one/two PYL1-like ABA receptors were preferentially expressed, whereas gene expression of the rest was low or almost undetectable in leaves under the following conditions: well-watered, heat, drought, heat+drought, salt or ABA treatment (Fig. 2A, B). Pd15 and Pd93 are PYL1-like receptors that show high nucleotide sequence identity and some of the 100 bp RNA-seq data were ambiguous reads, therefore we could not unequivocally assign them. Abiotic stress and ABA treatment did not substantially affect transcript expression of ABA receptors (Fig. 2A, B).

**Fig. 2.**
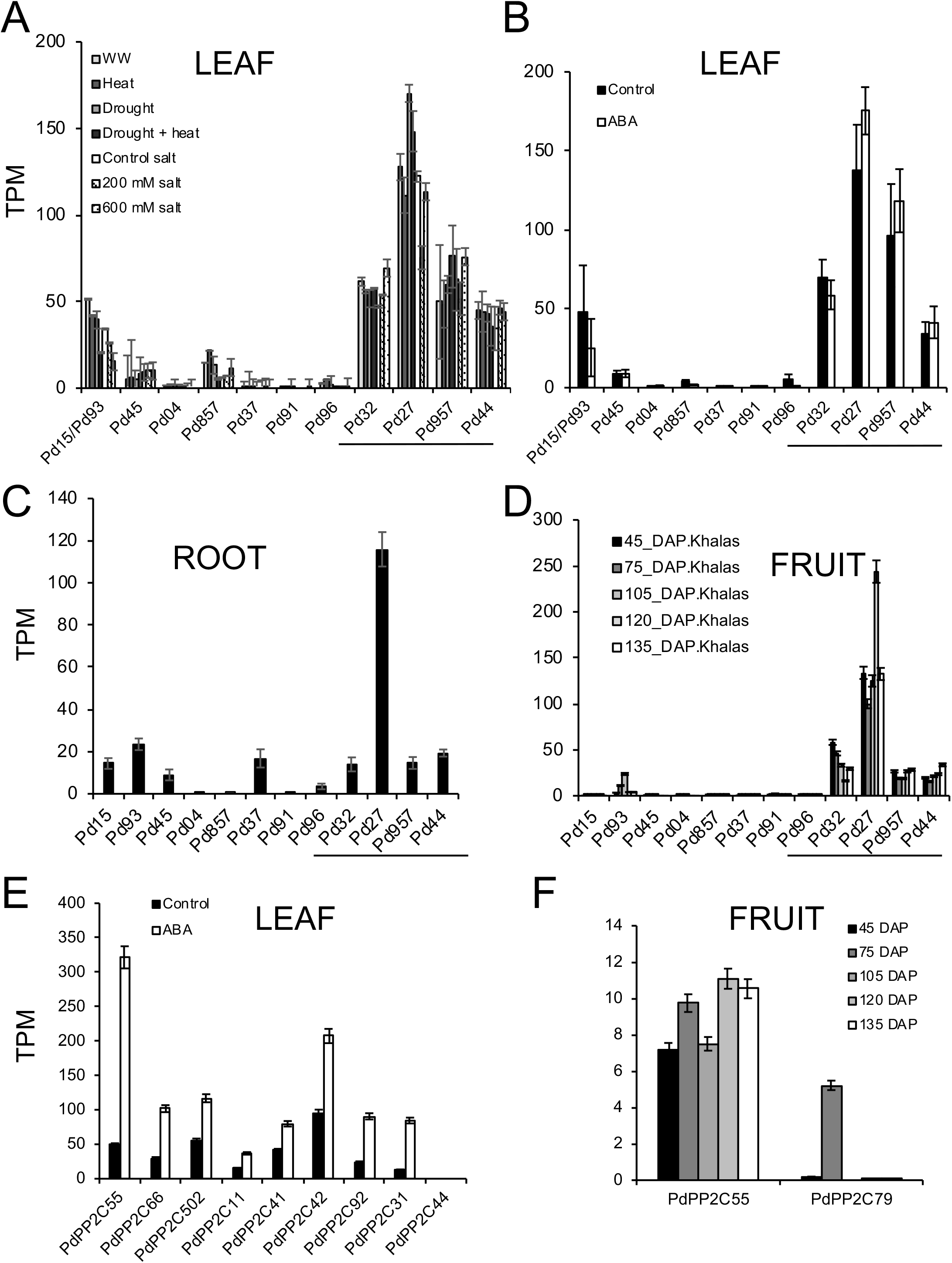
Relative gene expression of date palm receptors in abiotic stress conditions and different tissues was determined by RNA-Seq analysis. PYL8-like receptors are underlined. TPM (transcripts per million) normalizes transcript expression according to transcript length and sequencing depth. (A) Expression of Pd ABA receptors in leaves under well-watered (WW), heat, drought, heat+drought or salt treatment. (B) Expression of Pd ABA receptors in mock- or ABA-treated leaves. Data were obtained from the ArrayExpress archive (E-MTAB-5261) at EMBL-EBI. (C) Expression of Pd ABA receptors in root. Data were obtained from the Sequence Read Archive (experiment PRJNA497070) (Xiao et al. 2019). (D) Expression of Pd ABA receptors in fruit. Data were obtained from the Sequence Read Archive (experiment SRX122676) and correspond to different stages of fruit development after pollination: 45, 75, 105, 120 and 135 days after pollination (DAP) (Hazzouri et al., 2019). (E) Expression of Pd PP2Cs in mock- or ABA-treated leaves. Data were obtained from the ArrayExpress archive (E-MTAB-5261) at EMBL-EBI. (F) Expression of PdPP2C55 and PdPP2C79 in fruit at the indicated DAP (Hazzouri et al., 2019).

Given that date palm has a well-designed and specialized root system to optimally manage water uptake, we investigated expression of Pd ABA receptors in root tissue (Xiao et al., 2019). Expression of several receptors was detected in roots and particularly the PYL8-like Pd27 receptor was expressed at 5-6 fold higher levels than other receptors (Fig. 2C). PYL8 plays an important and non-redundant role for ABA signaling in *Arabidopsis* root (Antoni et al., 2013; Belda-Palazon et al., 2018). The predominant expression of Pd27 (and other PYL8-like receptors) in root tissue suggests that ABA signaling mediated by PYL8-like receptors might play a key role to maintain primary root growth at low water potential, to regulate root system architecture or for root hydrotropism, as it has been described in other plant species (Sharp et al., 2004; Dietrich et al., 2017; Orman-Ligeza et al., 2018). Expression of Pd15 and Pd93 could be distinguished in 150 bp RNA-seq reads of root tissue (Xiao et al., 2019), and also Pd45 or Pd37 were expressed in roots, although to 10-15 % levels compared to Pd27. Finally, RNA-seq data were also compiled for different fruit stages (at 45, 75, 105, 120 and 135 days after pollination, DAP) and again, PYL8-like ABA receptors showed the highest expression, whereas the rest were expressed at very low levels or undetectable (Fig. 2D).

### In vitro activity of Pd ABA receptors against PP2Cs of Arabidopsis and Phoenix dactylifera

Inhibition of clade A PP2Cs by ABA receptors is the key signature of their biochemical function (Park et al., 2009; Ma et al., 2009; Santiago et al., 2009). In order to biochemically characterize Pd ABA receptors we have obtained recombinant protein for five of them. We ordered synthetic genes of Pd15, Pd27, Pd32, Pd44 and Pd957, cloned them in pCR8 and next in pETM11 vector for expression in *E. coli*. Recombinant His-tagged protein was obtained for each receptor (Fig. 3A, top) and in addition to SDS-PAGE we performed native gel electrophoresis in order to test the oligomeric state of each protein (Fig. 3A, bottom). The PYL1-like Pd15 receptor migrated as a dimer, whereas the PYL8-like Pd27, 32, 44 and 957 receptors migrated mostly as monomers, which shows a good correspondence with the oligomeric state of the *Arabidopsis* relatives (Dupeux et al., 2011; Hao et al., 2011). Next, we measured the capability of Pd receptors to inhibit *Arabidopsis* phosphatase HAB1 using pNPP as a substrate. As a result, all Pd ABA receptors showed ABA-dependent inhibition of HAB1 (Fig. 3B).

**Fig. 3.**
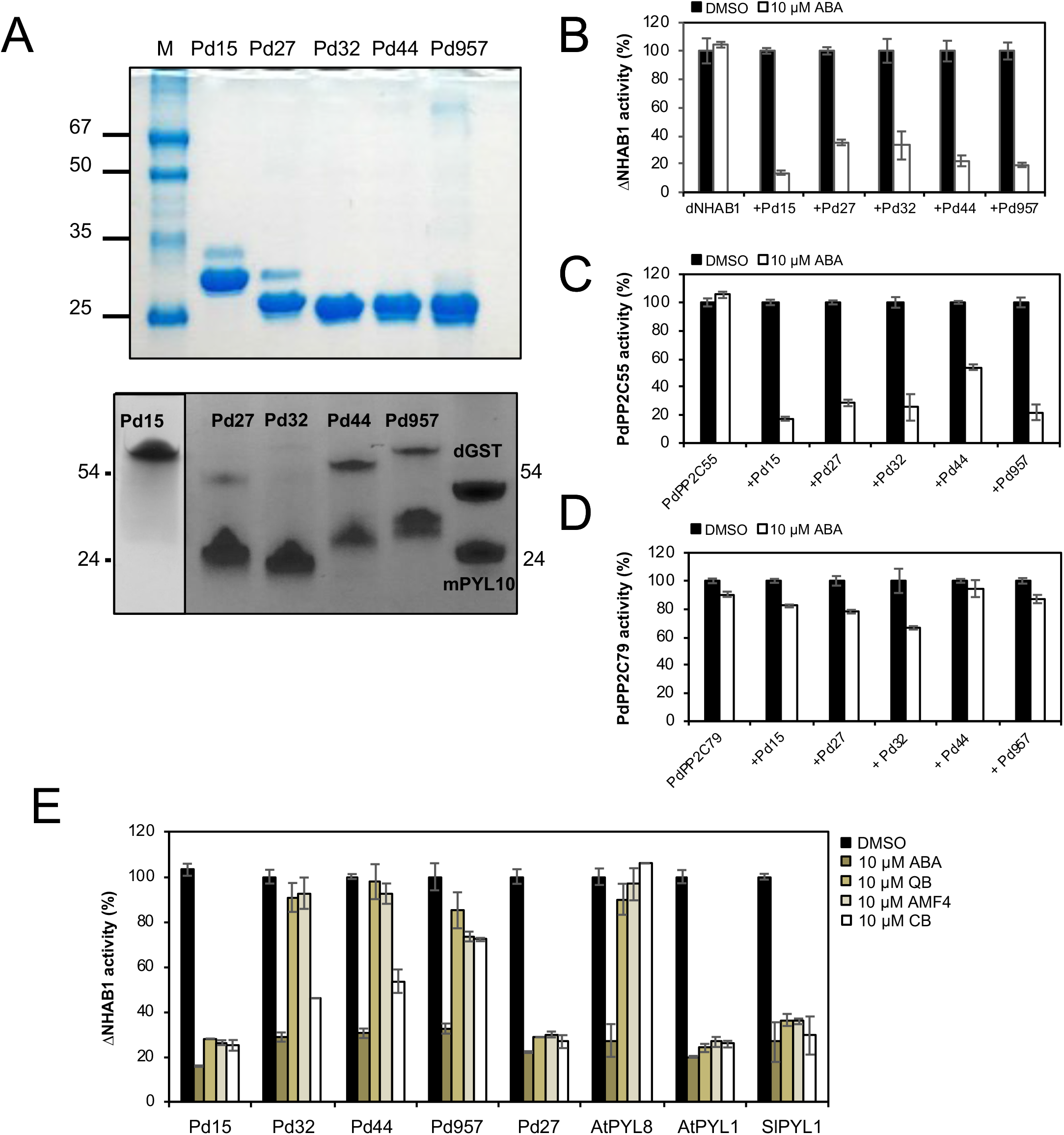
Expression, purification and analysis of recombinant Pd ABA receptors. (A) SDS-PAGE (top) and native gel electrophoresis (bottom) of Pd15, Pd27, Pd32, Pd44 and Pd957 ABA receptors. Molecular mass markers in the SDS PAGE are indicated in kDa, whereas in native gel we used monomeric PYL10 and dimeric GST as markers. (B, C, D) ABA-dependent PP2C inhibition mediated by Pd ABA receptors. PP2C activity was measured in vitro using the pNPP substrate (for ΔNHAB1) or the RRA(phosphoT)VA phosphopeptide (for PdPP2C55 and PdPP2C79) in the absence (DMSO) or presence of ABA at a 1:2 ratio (phosphatase:receptor). Data are averages ±SD for three independent experiments. Phosphatase activity in the absence of receptor and ABA was taken as 100 % activity. (E) Inhibition of ΔNHAB1 phosphatase activity in the presence of Pd, AtPYL8, AtPYL1 or SlPYL1 receptors mediated by 10 μM ABA or different ABA agonists: cyanabactin (CB), quinabactin (QB) and AMF4.

In order to identify Pd PP2Cs that might be a target of Pd ABA receptors, we performed a BLAST search in the date palm genome using ABI1 as a query to identify closely related Pd PP2Cs (Supplementary Fig. S1). *Arabidopsis* PP2Cs form a large family composed of 76 members (Schweighofer et al., 2004) and more than 100 genes in date palm show the PP2C signature (https://www.genome.jp/kegg-bin/show_organism?org=pda). We selected the top 10 PP2Cs that showed the highest sequence similarity to the catalytic PP2C domain of ABI1 as putative clade A PP2Cs from *Phoenix dactylifera* and we used the last two digits of the full KEGG code to name them (Supplementary Fig. S1). Further phylogenetic analysis revealed that some of these PP2Cs showed closer relationship to HAB1, AHG1 or PP2CA (Supplementary Fig. S1) Transcription profiles of PdPP2Cs were obtained from RNAseq studies performed by Müller et al., (2017). ABA treatment strongly upregulated expression of PdPP2Cs as occurs in *Arabidopsis* and other plant species (Fig. 2E) (Gonzalez-Guzman et al., 2014; Wang et al., 2019). We ordered synthetic genes for PdPP2C55 (the most expressed PP2C after ABA treatment), and PdPP2C79, which is only expressed during fruit development (Yin et al., 2012; Hazzouri et al., 2019) (Fig. 2F). PP2C79 shows high similarity with the *Arabidopsis* seed-specific PP2C AHG1, and a closely related PP2C is present in the oil palm (*Elaeis guineensis*) as well as other crops (Supplementary Fig. S2A and 2B). AHG1 plays an important role to regulate seed dormancy in *Arabidopsis* via interaction with DOG1; however, AHG1-like PP2Cs in monocot species have not been investigated yet (Née et al., 2017; Nishimura et al., 2018). The corresponding synthetic genes were cloned in pETM11 vector for expression in *E. coli* and recombinant His-tagged protein was obtained for both PP2Cs. PdPP2C55 phosphatase activity was measured using the RRA(phosphoT)VA phosphopeptide as a substrate because pNPP was a poor substrate for this phosphatase in contrast to HAB1. We found that all Pd ABA receptors showed ABA-dependent inhibition of PdPP2C55 (Fig. 3C). Therefore, Pd ABA receptors are functional in vitro to inhibit both HAB1 and PdPP2C55 in an ABA-dependent manner.

PdPP2C79, as previously reported for AHG1, lacks a conserved Trp residue required for efficient binding to ABA’s ketone group and *Arabidopsis* ABA receptors (Dupeux et al., 2011) (Supplementary Fig. S2A). In contrast, this Trp residue is conserved in other clade A PP2Cs (Trp385 in HAB1, Trp300 in ABI1) and establishes a water-mediated hydrogen-bond network (known as Trp lock) with the ketone group of ABA and the gate/latch loops of the receptor (Dupeux et al., 2011). Therefore, mutations in this conserved Trp residue (HAB1^W385A^ and ABI1^W300A^), or changes in neighboring residues as occurs in *Striga hermonthica* PP2C1, render receptor-insensitive PP2Cs that lead to attenuation of ABA signaling (Dupeux et al., 2011; Fujioka et al., 2019). In general, PP2C79 was quite refractory to inhibition by Pd ABA receptors and in the best case, Pd32 rendered only 30% inhibition of this phosphatase whereas inhibited by 80% PdPP2C55 at 10 μM ABA (Fig. 3D). Expression of PP2C79 in *Arabidopsis* led to plants that were strongly insensitive to ABA in germination and root growth assays (Supplementary Fig. 2C, D). Additionally, these plants showed higher water loss than wild-type Col-0 and were prone to wilting when watering was withheld (Supplementary Fig. 2E). Ectopic expression of PdPP2C79 in root tissue led to reduced sensitivity to ABA-mediated inhibition of root growth (Supplementary Fig. 2F). Given that in date palm PdPP2C79 is only expressed during fruit development, these results suggest that PdPP2C79 might provide a mechanism to attenuate ABA signaling at that stage in order to prevent developmental arrest induced by ABA accumulation (Xiao et al., 2019).

### Sensitivity of Pd ABA receptors to ABA agonists

ABA receptor agonists are a promising tool to control plant transpiration and drought resistance, and several agonists have been found to be active against *Arabidopsis* ABA receptors (Helander et al., 2016; Okamoto and Cutler, 2018). However, few examples have been reported of agonist activity against crop ABA receptors (Gonzalez-Guzman et al., 2014; Vaidya et al., 2019). We used Pd receptors as targets to test the effect of several ABA receptor agonists: quinabactin (QB) (Okamoto, 2013; Cao et al., 2013), AMF4 (Cao et al., 2017), and cyanabactin (CB) (Vaidya et al., 2017) (Fig. 3E). AtPYL1 and SlPYL1 were sensitive to ABA agonists that target dimeric receptors such as QB, CB and AMF4, and Pd15 behaved similarly, which further confirms that Pd15 belongs to the PYL1-like branch. Pd32 and Pd44 were resistant to QB and AMF4, but were sensitive to CB, which not only targets the dimeric PYR1 and PYL1 receptors in *Arabidopsis*, but also inhibits PYL5 efficiently (Vaidya et al., 2017). Pd957 sensitivity to QB, CB and AMF4 was low (only 20-30% inhibition of phosphatase activity) and AtPYL8 was even less sensitive to these agonists (Fig. 3E). Unexpectedly, Pd27 inhibited by 80% the activity of HAB1 in the presence of 10 μM of any agonist, which suggests that crop receptors can show different sensitivity to ABA agonists than *Arabidopsis* receptors. Vaidya et al. (2019) reported that a single amino acid difference between wheat PYL8 and Arabidopsis PYL8 led to high PP2C inhibitory effect of opabactin with wheat PYL8 and low effect with AtPYL8. Inspection of wheat subfamily I ligand-binding pockets revealed that the bulky Leu163 of AtPYL8 was replaced by a smaller Val179 in TaPYL8 (Vaidya et al. 2019). The Leu of AtPYL8 would cause steric clash for the binding of opabactin, whereas it seems that the smaller Val of TaPYL8 avoids it and leads to high agonist potency for PP2C inhibition. Similar minor changes occur in the sequence of Pd27 (or TaPYL8) compared to AtPYL8, for example Leu126 of AtPYL8 (close to the conserved latch loop) is replaced by Val143 in Pd27 and Val142 in TaPYL8 (Fig. 1B). Further structural work with crop receptors is required for a full understanding of agonist sensitivity.

### In vivo activity of Pd ABA receptors

We tested the in vivo activity of Pd ABA receptors by measuring ABA signaling (LUC expression driven by the ABA-responsive RD29B promoter) induced in protoplasts upon ABA perception. To this end we transfected effector DNA constructs together with an ABA-responsive luciferase reporter in the absence or presence of exoqenous ABA (Fujii et al., 2009). A constitutively expressed GUS reporter was used for normalization and accordingly the LUC/GUS activity ratio was used to measure ABA signaling (Fujii et al., 2009). In the presence of ABA, LUC activity was elicited by the effector ABF2 construct, whereas co-expression of HAB1 abolished ABA-induced LUC activity (Fig. 4A). Introduction of the PYL8 receptor together with the ABF2+HAB1 expression cassettes restored ABA signaling as well as introduction of each Pd ABA receptor. In the case of Pd27 and Pd957, in the absence of exogenous ABA addition, some induction of LUC activity was also observed. In order to visualize the subcellular localization of Pd ABA receptors and PdPP2Cs, we performed transient expression in *Nicotiana benthamiana*. In addition to GFP fusion proteins of receptors or phosphatases, we also co-expressed the nucleolar marker fibrillarin-RFP (Belda-Palazon et al., 2019). In the case of PdPP2C55 and PdPP2C79, both PP2Cs were predominantly localized in the nucleus of *N. benthamiana* epidermal leaf cells (Fig. 4B). We found that Pd ABA receptors were localized both in nucleus and cytosol; however, Pd27 and Pd957 showed preferential nuclear localization (Fig. 4C). Expression of the GFP fusion proteins was verified by immunoblot analysis (Fig. 4D).

**Fig. 4.**
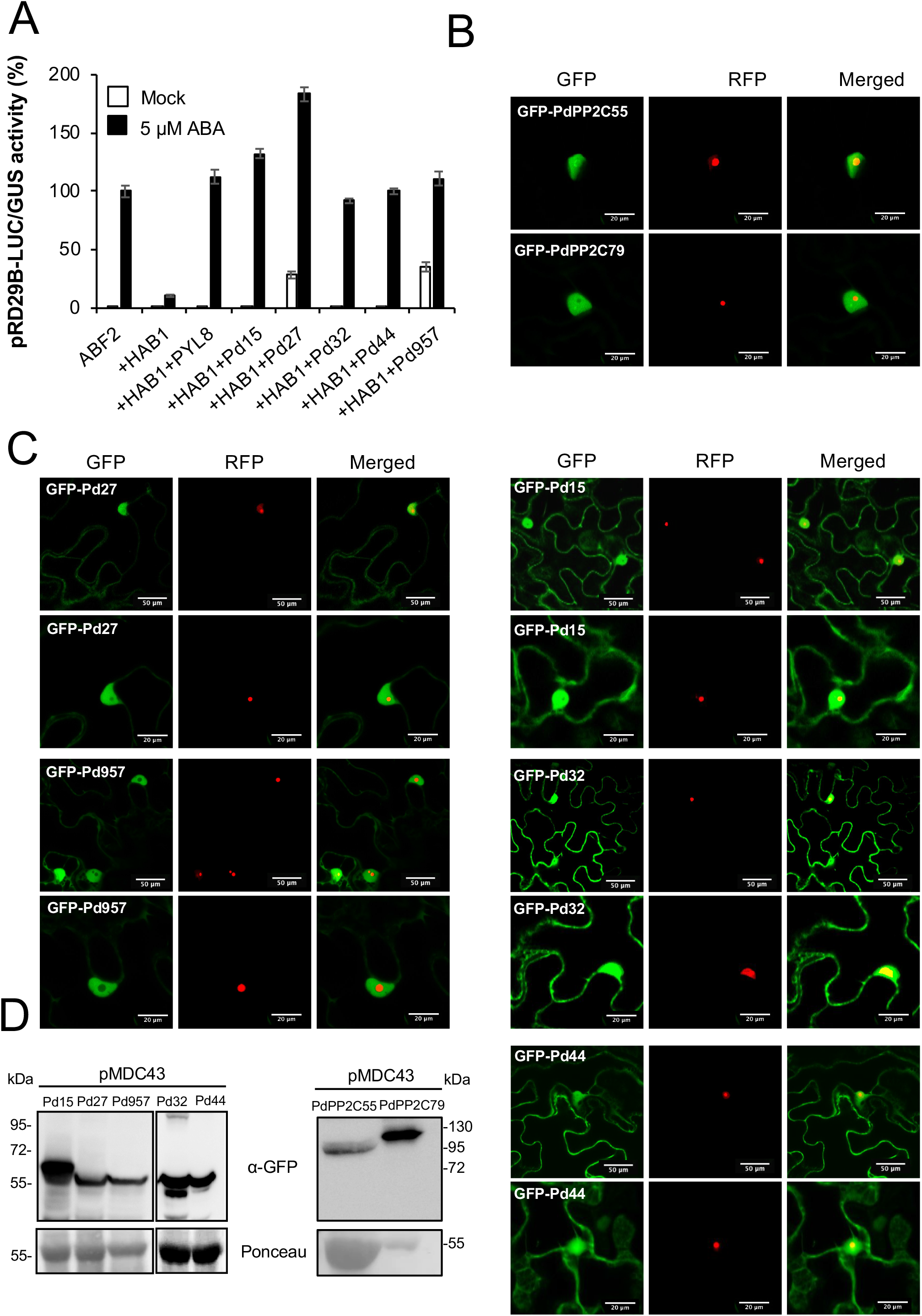
ABA signaling mediated by Pd ABA receptors in protoplasts and subcellular localization of PdPP2Cs and ABA receptors. (A) Protoplasts of wildtype Col-0 were transfected with plasmids encoding the indicated proteins in absence or presence of ABA. LUC activity was measured in protein extracts prepared from protoplast suspensions 6 h after transfection, either in the absence or presence of 5 μM exogenous ABA (added 3 h after transfection). The activity of the LUC reporter was normalized with GUS activity and the ratio LUC/GUS is indicated. (B) (C) Subcellular localization of PdPP2Cs and ABA receptors. PdPP2C55 and PdPP2C79 localize predominantly to nucleus, whereas Pd ABA receptors localize to nucleus and cytosol upon transient expression in *N. benthamiana* leaf cells. Confocal images of transiently transformed epidermal cells expressing GFP-PdPP2C55, GFP-PdPP2C79, GFP-Pd15, GFP-Pd27, GFP-Pd32, GFP-Pd44 or GFP-Pd957. The GFP channel shows the subcellular localization of the GFP fusion proteins, whereas the RFP channel shows the nucleolar marker fibrillarin-RFP. Merged indicates the overlap of the GFP and RFP channels. Scale bars correspond to 50 or 20 μm. (D) Immunoblotting analysis using anti-GFP antibodies was used to verify the expression of the corresponding fusion proteins.

Overexpression of monomeric ABA receptors confers enhanced response to ABA and drought resistance (Santiago et al., 2009; Gonzalez-Guzman et al., 2014; Yang et al., 2016, Mega et al., 2019), and tomato ABA receptors are functional when expressed in *Arabidopsis* (Gonzalez-Guzman et al., 2014). We were interested in testing whether monocot ABA receptors such as date palm receptors were functional in a dicot species as *Arabidopsis*. We expressed GFP-tagged Pd ABA receptors in *Arabidopsis* as a first step towards analyzing their role in ABA signaling. Expression of the proteins in vegetative tissue was analyzed by immunoblot analysis and since ABA promotes stabilization and accumulation of the *Arabidopsis* PYL8 receptor (Belda-Palazon et al., 2018), we analyzed protein expression in mock- and ABA-treated plants (Fig. 5A). Pd15 protein level did not substantially change in mock- and ABA-treated samples, whereas PYL8-like Pd receptors showed ABA-induced accumulation (Fig. 5A). Therefore, this result reveals that ABA-induced accumulation of PYL8-like receptors also occurs when monocot receptors are expressed in *Arabidopsis* (Belda-Palazon et al., 2018; Fig. 5A). Moreover, analysis of ABA response in transgenic lines overexpressing GFP-Pd receptors, revealed enhanced sensitivity to ABA mediated inhibition of seedling establishment compared to wild-type Col-0, indicating that Pd receptors are functional when expressed in *Arabidopsis* (Fig. 5B). In agreement with the Pd27 sensitivity to ABA agonists (Fig. 3E), we observed enhanced sensitivity to QB and AMF4 in plants overexpressing GFP-Pd27 compared with wild-type Col-0 (Fig. 5C). Finally, we confirmed ABA-induced accumulation of the GFP-Pd27 ABA receptor by CLSM imaging, both in root and shoot tissues (Fig. 5D, E). Likewise, GFP-Pd32, another PYL8-like ABA receptor, also accumulates in both tissues in response to ABA, whereas Pd15, which is a PYL1-like ABA receptor, does not accumulate in response to ABA (Fig. 5D, E).

**Fig. 5.**
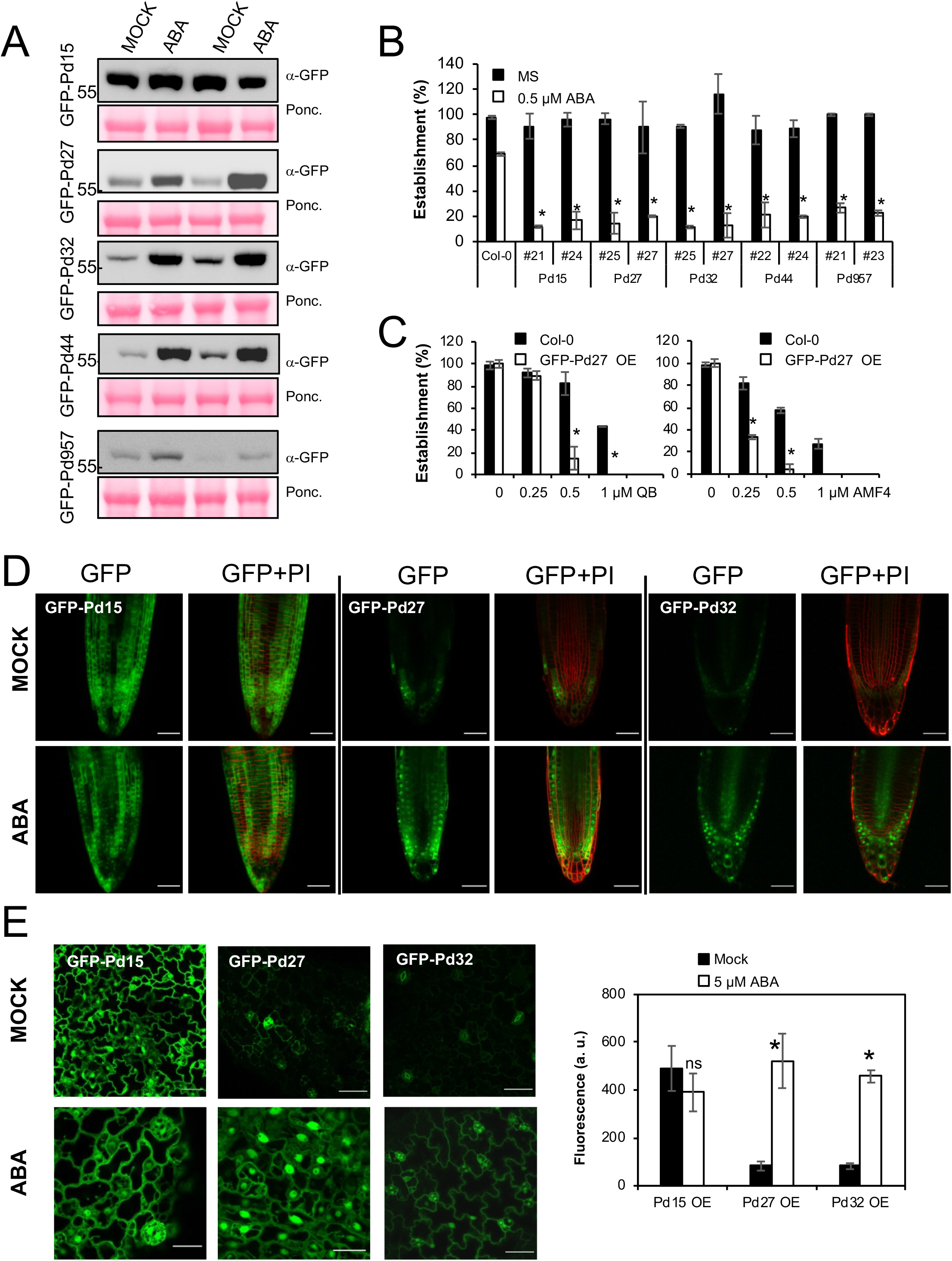
PdPYL8-like receptors expressed in *Arabidopsis* accumulate higher levels upon ABA-treatment. (A) Immunoblot analysis using anti-GFP antibody shows that ABA treatment enhances protein levels of PdPYL8-like receptors (two independent *Arabidopsis* T3 transgenic lines for each palm receptor). Ponceau staining is shown below. (B) GFP-Pd fusion proteins are functional in *Arabidopsis*. ABA-mediated inhibition of seedling establishment in transgenic lines compared with non-transformed Col-0 plants. * indicates P<0.05 (Student’s t test) when comparing data of transgenic lines to non-transformed Col-0 plants in the same assay conditions. Seedlings were scored for the presence of both green cotyledons and the first pair of true leaves 7 days after sowing. (C) Enhanced sensitivity to QB and AMF4 of GFP-Pd27 OE plants compared with wild-type Col-0. Approximately 25 seeds of the indicated genotype (three independent experiments) were sown on MS multi-well plates lacking or supplemented with the indicated concentration of QB or AMF4, and seedling establishment was scored for the presence of both green cotyledons and the first pair of true leaves after 7 d. Values are averaqes ±SD. The asterisk indicates p<0.01 (Student’s t test) with respect to wild type assayed in the same conditions. (D, E) GFP-Pd27 and GFP-Pd32 accumulate after ABA treatment, whereas GFP-Pd15 does not. CLSM imaging of *Arabidopsis* root apex (D) or leaf tissue (E) in lines expressing GFP-Pd15, GFP-Pd27 or GFP-Pd32 that were mock- or ABA-treated for 1 h. Two independent lines were analyzed and representative images obtained with one line are shown. Fluorescence was quantified (arbitrary units) in 10 leaves (three independent experiments) of each transgenic line using images acquired by CLSM.

### Overexpression of Pd27 confers drought resistance

We also expressed HA-tagged Pd ABA receptors in *Arabidopsis* and analyzed ABA response of the transgenic lines in seedling establishment and root growth assays (Fig. 6A-C). As GFP-Pd receptor lines, overexpression of HA-Pd ABA receptors conferred enhanced response to ABA compared to wild type. In order to test whether palm receptors can confer enhanced drought resistance, we selected transgenic lines overexpressing Pd27 because this receptor is the most expressed in all palm tissues analyzed (Fig. 2). Plants were grown under normal watering conditions for 2 weeks and then irrigation was stopped for 20 days. Non-transformed Col-0 plants wilted, in contrast to transgenic lines that express Pd27 (Fig. 6D). The survival of the plants was scored 5 days after watering was resumed and enhanced survival was found for transqenic plants expressing Pd27 (Fig. 6E). Transpiration was monitored using infrared thermography and we found that transgenic plants expressing Pd27 showed higher leaf temperature than wild-type Col-0 plants upon spraying with ABA (Fig. 7A). Further physiological analyses were conducted using normalized gravimetric water loss experiments to measure transpiration of wild-type Col-0 and Pd27 OE plants, which was recorded under different soil water potentials, ranging from well-watered (potential near to 0 MPa) to drought conditions (−2.5 MPa) (Fig. 7B). As a result, we found that Pd27 OE plants showed lower transpiration than wild type. To correlate transpiration and ABA levels, we measured endogenous ABA content (ng/g dry weight) by HPLC/MS analysis in the different conditions of soil water potential (SWP) assayed (Fig. 7C). Interestingly, Pd27 OE plants had a lower ABA content than wild type, which was particularly evident under drought conditions. Therefore, the combination of transpiration data and ABA content reveals that Pd27 OE plants were more efficient to reduce transpiration at each ABA concentration, particularly when ABA levels increase (Fig. 7D).

**Fig. 6.**
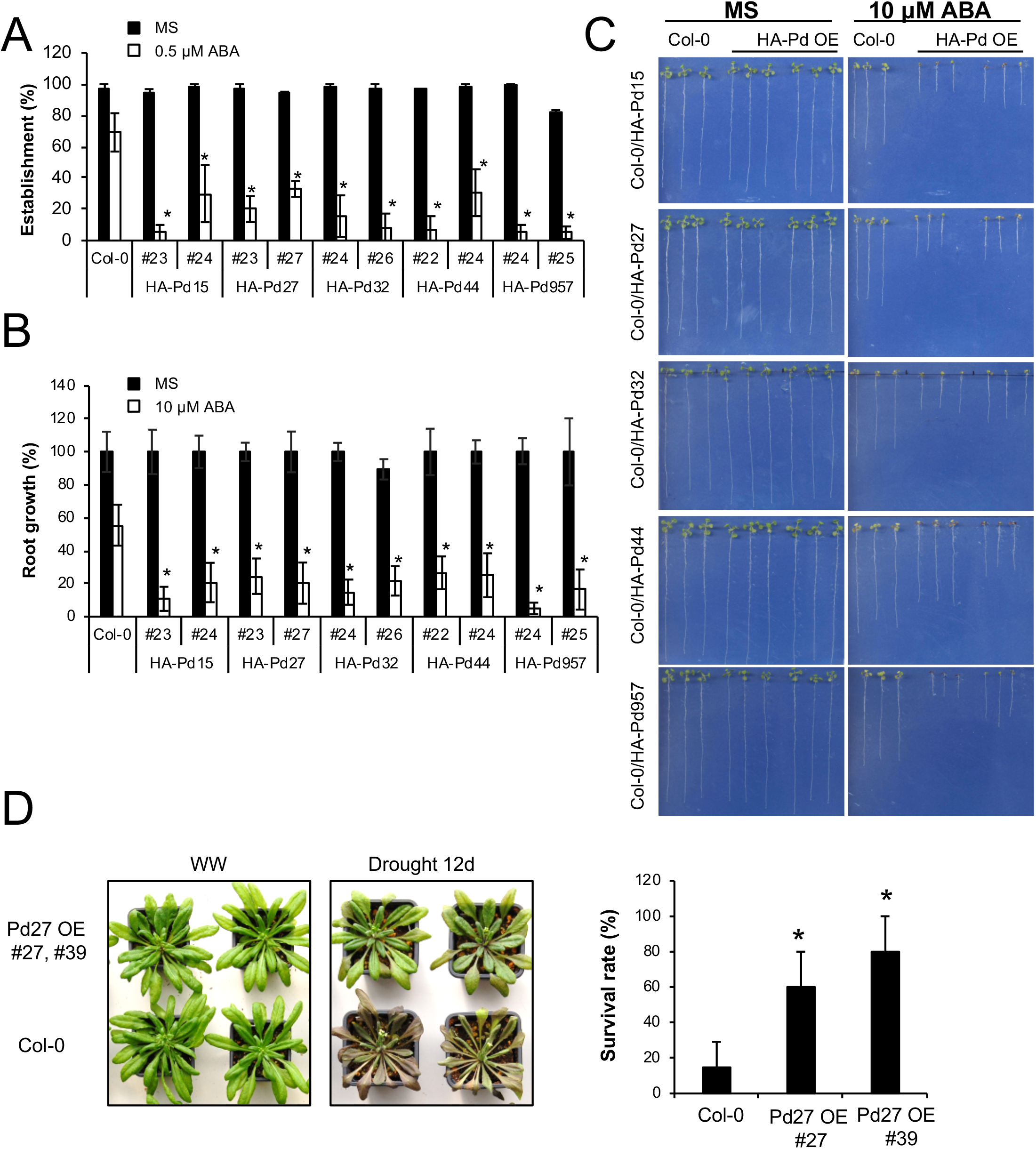
Enhanced sensitivity to ABA conferred by overexpression of Pd ABA receptors. (A) Enhanced sensitivity to ABA-mediated inhibition of seedling establishment. Approximately 100 seeds of each genotype (three independent experiments) were sown on MS plates lacking or supplemented with 0.5 μM ABA and scored for the presence of both green cotyledons and the first pair of true leaves after 7 d. Values are averages ±SD. The asterisk indicates p<0.01 (Student’s t test) with respect to wild type assayed in the same conditions. (B) Enhanced sensitivity to ABA-mediated inhibition of root growth. Quantification of ABA-mediated inhibition of root growth in the indicated genotypes compared to wild type. The asterisk indicates p<0.01 (Student’s t test) with respect to wild type assayed in the same conditions. (C) The photoqraphs show representative seedlings 10 d after the transfer of 4-d-old seedlinqs to MS plates lacking or supplemented with 10 μM ABA. (D) Enhanced drought resistance of Pd27 OE plants compared to Col-0. Irrigation was withdrawn in three-week-old plants for 20 d and after rewatering, survival of the plants was scored after 5 d. Data are means of three independent experiments ±SD (n=10 per experiment). The survival percentage 5 d after rewatering is indicated in histograms.

**Fig. 7.**
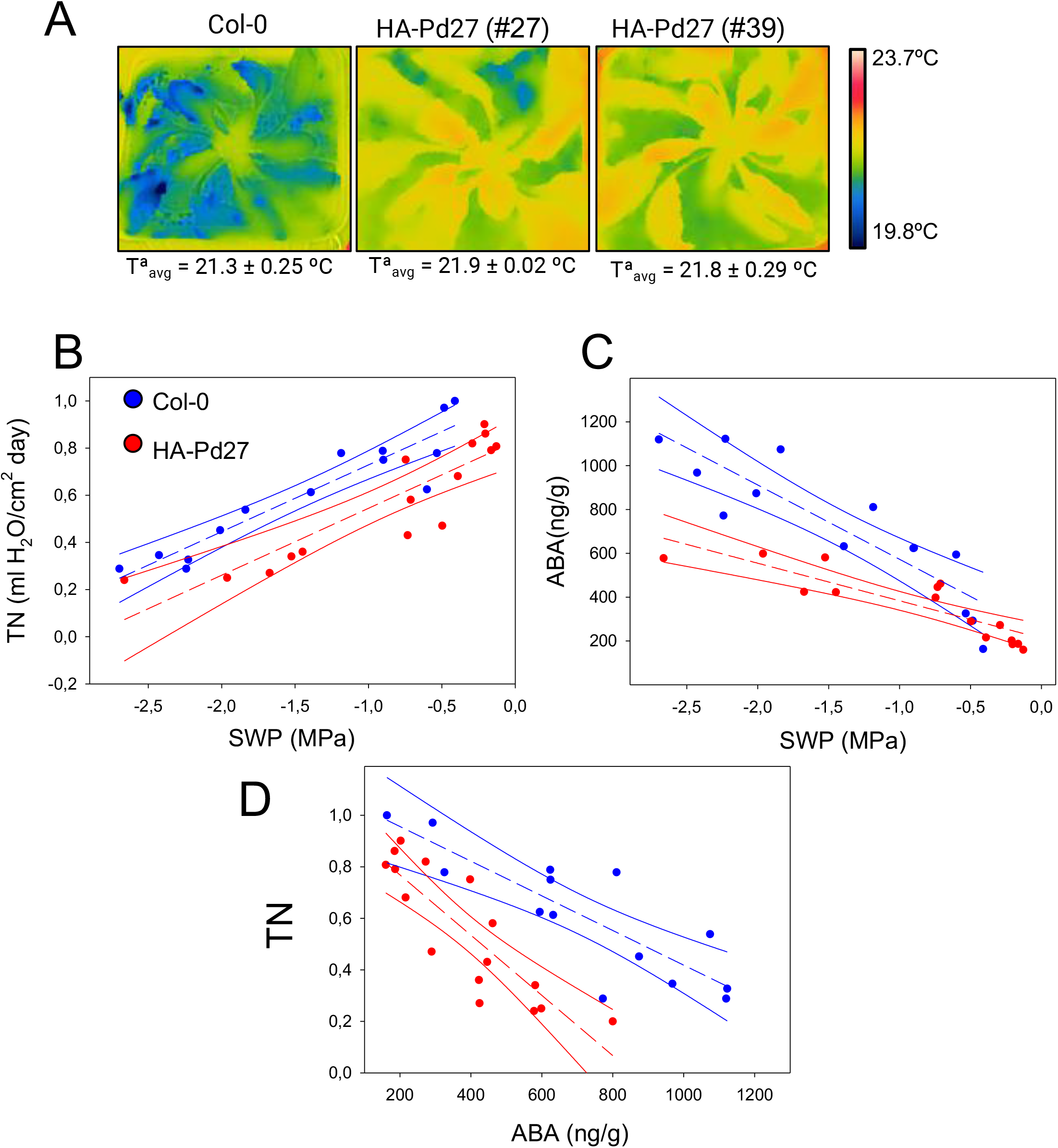
Plants OE Pd27 show lower transpiration than wild type. (A) Infrared thermography analysis shows higher leaf temperature of two independent *Arabidopsis* transgenic lines OE Pd27. False-color images of wild-type Col-0 and Pd27 OE plants (lines #27 and #39) representing leaf temperature, which was quantified by infrared thermal imaging. Data are means ±SD (n=5, aprox. 1000 measurements of square pixels from multiple leaves of each plant). (B) Transpiration assay under different soil water potential (SWP) conditions. (C) ABA quantification under different SWP conditions. (D) ABA lowers transpiration more efficiently in plants OE Pd27 than in wild type. For experiments of B, C and D panels, n=16. In B, C and D differences in transpiration (TN), endogenous ABA and transpiration (TN), respectively, were significant due to genotype (p=0.001, p=0.003 and p= 0.001, respectively) after ANOVA using SWP (B, C) and ABA (D) as covariant.

## Discussion

Abiotic stress is challenging plant agriculture and ABA is a key phytohormone to trigger plant adaptation to stressors, particularly under drought conditions (Yang et al., 2019). Date palm is very well adapted to heat and drought conditions, but little is known about the underlying molecular mechanisms (Arab et al., 2016). Guard cell ABA signaling mediated by PdOST1 limits transpiration water loss in date palm, presumably through ABA-dependent activation mediated by ABA receptors (Müller et al., 2017). Recent findings have revealed the role of certain *Arabidopsis* receptors in stomatal closure signal integration, and for instance PYL2 is critical for guard cell ABA-induced responses whereas PYL4 and PYL5 are essential in the responses to CO2 (Dittrich et al., 2019). In this work we have identified which members of the date palm family of ABA receptors are particularly relevant for abiotic stress response. The phylogenetic analysis revealed that PdPYLs can be divided into three subfamilies, which correspond to PYL1-like, PYL4-like and PYL8-like *Arabidopsis* relatives. Based on RNA seq analyses of PdPYLs, and somehow unexpectedly, we discovered that PYL8-like receptors play a major role for palm response to abiotic stress. This result contrasts with *Arabidopsis* ABA receptors, where members of the three subfamilies are significantly expressed in different tissues and growth conditions, and particularly *Arabidopsis* dimeric receptors play an important role to regulate stomatal aperture and adaptation to drought (Gonzalez-Guzman et al., 2012; Okamoto et al., 2013; Dittrich et al., 2019). In contrast to *Arabidopsis*, four date palm PYL8-like ABA receptors were predominantly expressed under heat, salt, drought or osmotic stress conditions. In Arabidopsis, ABA treatment induced their accumulation, which was only previously observed for AtPYL8; therefore, it seems that date palm has expanded this adaptive mechanism to a full branch of ABA receptors for efficient response to abiotic stress.

Specifically, Pd27 was the most expressed receptor in all the conditions analyzed, including root and fruit tissues. Analysis of transgenic palms is not feasible; therefore, we investigated Pd27 in *Arabidopsis* lines. We measured the effect of Pd27 on transpiration under different soil water potentials, which range from well-watered to drought conditions. In all the conditions tested, expression of Pd27 led to lower transpiration than wild type, and interestingly this effect was accompanied by a reduction in ABA levels, particularly at negative soil water potentials (Fig. 7B and 7C). As a result, reduction of transpiration in Pd27 OE plants per ng ABA was more efficient than in wild type (Fig. 7D). Drought leads to 10-40 fold increase of ABA levels (Cutler et al., 2010); therefore we suggest that stabilization of Pd27 (and likely other PYL8-like receptors such as Pd32) induced by the hormone increase might boost ABA signaling, because higher levels of the receptor will accumulate when ABA levels increase. Transcriptional and posttranslational mechanisms limit accumulation of ABA receptors in *Arabidopsis* (Fernandez et al., 2020); but transcriptional downrequlation either by ABA or different stressors was not observed in palm receptors. Moreover, ABA treatment in *Arabidopsis* transgenic lines led to higher accumulation of palm receptors. If this feature of the PYL8-like family is conserved in other crops adapted to drought, it might represent a valuable trait to be introduced in crops lacking such adaptation. Pd27 enhanced drought tolerance upon expression in *Arabidopsis* and was able to bind different ABA agonists, whose application together with Pd27 overexpression might enhance even further plant response to ABA. The result of combining Pd27 OE plants with spraying of different agonists when stress occurs might represent a powerful tool for tuning water use dynamically (Cao et al., 2017; Dejonghe et al., 2018; Vaidya et al., 2019). Additionally, the chemical approach benefits from the higher expression of the ABA receptor because lower dosage of the chemical is needed.

This study also reports the ABA- and receptor-resistant activity of the phosphatase PdPP2C79, as well as the extended presence of AHG1-like PP2Cs in different crops, bot monocot and dicot species. Given that AHG1 is a target of a key QTL that regulates seed dormancy such as *Arabidopsis* DOG1 (Nee et al., 2017; Nishimura et al., 2018), these results suggest that AHG1-DOG1 interaction might be evolutionary conserved and deserves further research in crops. Additionally, loss of the Trp lock in AHG1-like PP2Cs seems to have markedly reduced their regulation by ABA and ABA receptors, which likely enables AHG1 activity at plant developmental stages where high ABA concentration accumulates.

## Supplementary data

Supplementary data is available at JXB online

**Supplementary Fig. S1.** Phylogenetic analysis reveals the presence of HAB1-like, AHG1-like and PP2CA-like PP2Cs in date palm

**Supplementary Fig. S2.** AHG1-like PP2Cs lacking the Trp lock residue are present both in monocot and dicot plant species.

**Supplementary Fig. S3.** Expression of PdPP2C79 in Arabidopsis reduces ABA sensitivity.

**Supplementary Table S1.** List of antibodies and oligonucleotides used in this work.

## Data Availability

All data supporting the findings of this study are available within the article and its supplementary materials published online. RNA seq crude data are available in the Sequence Read Archive.

## Acknowledgements

This study was funded by the Distinguished Scientist Fellowship Research Program (DSFP) of King Saud University to P.L.R. The authors JJ., RR-P and IG-M were supported by a research contract funded by the DSFP.

## Author contribution

All authors designed and supervised experiments; IGM, AC, JL-J, MM, CO, JJ, RR-P, GP, BB-P performed experimental analyses; PLR devised the project and wrote the manuscript with contributions of CO and MM; and all authors analyzed the data and approved the final text

## Notes

### Competing Interest Statement

The authors have declared no competing interest.

## References

Al-Dous EK, George B, Al-Mahmoud ME, et al. 2011. De novo genome sequencing and comparative genomics of date palm (*Phoenix dactylifera*). Nature Biotechnology 29, 521–527.

Al-Harrasi I, Al-Yahyai R, Yaish MW. 2018. Differential DNA methylation and transcription profiles in date palm roots exposed to salinity. PLoS ONE 13, e0191492.

Al-Mssallem IS, Hu S, Zhang X, et al. 2013. Genome sequence of the date palm *Phoenix dactylifera* L. Nature Communications 4, 2274.

Antoni R, Gonzalez-Guzman M, Rodriguez L, et al. 2013. PYRABACTIN RESISTANCE1-LIKE8 plays an important role for the regulation of abscisic acid signaling in root. Plant Physiology 161, 931–941.

Antoni R, Gonzalez-Guzman M, Rodriguez L, Rodrigues A, Pizzio GA, Rodriguez PL. 2012. Selective inhibition of Clade A phosphatases type 2C by PYR/PYL/RCAR abscisic acid receptors. Plant Physiology 158, 970–980.

Arab L, Kreuzwieser J, Kruse J, Zimmer I, Ache P, Alfarraj S, Al-Rasheid KAS, Schnitzler JP, Hedrich R, Rennenberg H. 2016. Acclimation to heat and drought-Lessons to learn from the date palm (*Phoenix dactylifera*). Environmental and Experimental Botany 125, 20–30.

Arbona V, Zandalinas SI, Manzi M, González-Guzmán M, Rodriguez PL, Gómez-Cadenas A. 2017. Depletion of abscisic acid levels in roots of flooded Carrizo citrange (*Poncirus trifoliata* L. Raf. × *Citrus sinensis* L. Osb.) plants is a stress-specific response associated to the differential expression of PYR/PYL/RCAR receptors. Plant Molecular Biology 93, 623–640.

Baykov AA, Evtushenko OA, Avaeva SM. 1988. A malachite green procedure for orthophosphate determination and its use in alkaline phosphatase-based enzyme immunoassay. Analytical Biochemistry 171, 266–270.

Belda-Palazon B, Gonzalez-Garcia MP, Lozano-Juste J, et al. 2018. PYL8 mediates ABA perception in the root through non-cell-autonomous and ligand-stabilization-based mechanisms. Proceedings of the National Academy of Sciences of the United States of America 115, E11857–E11863.

Belda-Palazon B, Julian J, Coego A, et al. 2019. ABA inhibits myristoylation and induces shuttling of the RGLG1 E3 ligase to promote nuclear degradation of PP2CA. Plant Journal 98, 813–825.

Bourgis F, Kilaru A, Cao X, Ngando-Ebongue GF, Drira N, Ohlrogge JB, Arondel V. 2011. Comparative transcriptome and metabolite analysis of oil palm and date palm mesocarp that differ dramatically in carbon partitioning. Proceedings of the National Academy of Sciences of the United States of America 108, 12527–12532.

Cao M, Liu X, Zhang Y, et al. 2013. An ABA-mimicking ligand that reduces water loss and promotes drought resistance in plants. Cell Research 23, 1043–1054.

Cao MJ, Zhang YL, Liu X, et al. 2017. Combining chemical and genetic approaches to increase drought resistance in plants. Nature Communications 8, 1183.

Christmann A, Hoffmann T, Teplova I, Grill E, Müller A. 2005. Generation of active pools of abscisic acid revealed by in vivo imaging of water-stressed arabidopsis. Plant Physiology 137, 209–219.

Clough SJ, Bent AF. 1998. Floral dip: A simplified method for Agrobacterium-mediated transformation of *Arabidopsis thaliana*. Plant Journal 16, 735–743.

Cutler SR, Rodriguez PL, Finkelstein RR, Abrams SR. 2010. Abscisic Acid: Emergence of a Core Signaling Network. Annual Review of Plant Biology 61, 651–679.

Dejonghe W, Okamoto M, Cutler SR. 2018. Small molecule probes of ABA biosynthesis and signaling. Plant and Cell Physiology 59, 1490–1499.

De Ollas C, Segarra-Medina C, González-Guzmán M, Puertolas J, Gómez-Cadenas A. 2019. A customizable method to characterize *Arabidopsis thaliana* transpiration under drought conditions. Plant Methods 15.

DIetrich D, Pang L, Kobayashi A, et al. 2017. Root hydrotropism is controlled via a cortex-specific growth mechanism. Nature Plants 3, 17057.

Dittrich M, Mueller HM, Bauer H, et al. 2019. The role of Arabidopsis ABA receptors from the PYR/PYL/RCAR family in stomatal acclimation and closure signal integration. Nature Plants 5, 1002–1011.

Dupeux F, Antoni R, Betz K, et al. 2011. Modulation of abscisic acid signaling in vivo by an engineered receptor-insensitive protein phosphatase type 2C allele. Plant Physiology 156, 106–116.

Durgbanshi A, Arbona V, Pozo O, Miersch O, Sancho J V., Gómez-Cadenas A. 2005. Simultaneous determination of multiple phytohormones in plant extracts by liquid chromatography-electrospray tandem mass spectrometry. Journal of Agricultural and Food Chemistry 53, 8437–8442.

Fernandez MA, Belda-Palazon B, Julian J, Coego A, Lozano-Juste J, Iñigo S, Rodriguez L, Bueso E, Goossens A, Rodriguez PL. 2020. RBR-type E3 ligases and the ubiquitin-conjugating enzyme UBC26 regulate abscisic acid receptor levels and signaling. Plant Physiology 182, 1723–1742.

Fujii H, Chinnusamy V, Rodrigues A, Rubio S, Antoni R, Park SY, Cutler SR, Sheen J, Rodriguez PL, Zhu JK. 2009. In vitro reconstitution of an abscisic acid signalling pathway. Nature 462, 660–664.

Fujioka H, Samejima H, Suzuki H, Mizutani M, Okamoto M, Sugimoto Y. 2019. Aberrant protein phosphatase 2C leads to abscisic acid insensitivity and high transpiration in parasitic Striga. Nature Plants 5, 258–262.

Gonzalez-Guzman M, Pizzio GA, Antoni R, et al. 2012. Arabidopsis PYR/PYL/RCAR receptors play a major role in quantitative regulation of stomatal aperture and transcriptional response to abscisic acid. Plant Cell 24, 2483–2496.

Gonzalez-Guzman M, Rodriguez L, Lorenzo-Orts L, et al. 2014. Tomato PYR/PYL/RCAR abscisic acid receptors show high expression in root, differential sensitivity to the abscisic acid agonist quinabactin, and the capability to enhance plant drought resistance. Journal of Experimental Botany 65, 4451–4464.

Gros-Balthazard M, Hazzouri KM, Flowers JM. 2018. Genomic insights into date palm origins. Genes 9.

Hazzouri KM, Flowers JM, Visser HJ, et al. 2015. Whole genome re-sequencing of date palms yields insights into diversification of a fruit tree crop. Nature Communications 6, 8824.

Hazzouri KM, Gros-Balthazard M, Flowers JM, et al. 2019. Genome-wide association mapping of date palm fruit traits. Nature Communications 10, 4680.

He Y, Hao Q, Li W, Yan C, Yan N, Yin P. 2014. Identification and characterization of ABA receptors in oryza sativa. PLoS ONE 9, e95246.

He Z, Zhong J, Sun X, Wang B, Terzaghi W, Dai M. 2018. The maize ABA receptors ZmPYl8, 9, and 12 facilitate plant drought resistance. Frontiers in Plant Science 9, 422.

Helander JDM, Vaidya AS, Cutler SR. 2016. Chemical manipulation of plant water use. Bioorganic and Medicinal Chemistry 24, 493–500.

Ma Y, Szostkiewicz I, Korte A, Moes D, Yang Y, Christmann A, Grill E. 2009. Regulators of PP2C phosphatase activity function as abscisic acid sensors. Science 324, 1064–1068.

Mega R, Abe F, Kim JS, et al. 2019. Tuning water-use efficiency and drought tolerance in wheat using abscisic acid receptors. Nature Plants 5, 153–159.

Miao C, Xiao L, Hua K, Zou C, Zhao Y, Bressan RA, Zhu JK. 2018. Mutations in a subfamily of abscisic acid recepto genes promote rice growth and productivity. Proceedings of the National Academy of Sciences of the United States of America 115, 6058–6063.

Moes D, Himmelbach A, Korte A, Haberer G, Grill E. 2008. Nuclear localization of the mutant protein phosphatase abi1 is required for insensitivity towards ABA responses in Arabidopsis. Plant Journal 54, 806–819.

Müller HM, Schäfer N, Bauer H, et al. 2017. The desert plant *Phoenix dactylifera* closes stomata via nitrate-regulated SLAC1 anion channel. New Phytologist 216, 150–162.

Née G, Kramer K, Nakabayashi K, Yuan B, Xiang Y, Miatton E, Finkemeier I, Soppe WJJ. 2017. DELAY of GERMINATION1 requires PP2C phosphatases of the ABA signalling pathway to control seed dormancy Nature Communications 8.

Nishimura N, Tsuchiya W, Moresco JJ, et al. 2018. Control of seed dormancy and germination by DOG1-AHG1 PP2C phosphatase complex via binding to heme. Nature Communications 9.

Okamoto M, Cutler SR. 2018. Chemical control of ABA receptors to enable plant protection against water stress. Methods in Molecular Biology. 127–141.

Okamoto M, Peterson FC, Defries A, Park SY, Endo A, Nambara E, Volkman BF, Cutler SR. 2013. Activation of dimeric ABA receptors elicits guard cell closure, ABA-regulated gene expression, and drought tolerance. Proceedings of the National Academy of Sciences of the United States of America 110, 12132–12137.

Orman-Ligeza B, Morris EC, Parizot B, et al. 2018. The Xerobranching Response Represses Lateral Root Formation When Roots Are Not in Contact with Water. Current Biology 28, 3165–3173.e5.

Park SY, Fung P, Nishimura N, et al. 2009. Abscisic acid inhibits type 2C protein phosphatases via the PYR/PYL family of START proteins. Science 324, 1068–1071.

Planes MD, Niñoles R, Rubio L, et al. 2015. A mechanism of growth inhibition by abscisic acid in germinating seeds of *Arabidopsis thaliana* based on inhibition of plasma membrane H+-ATPase and decreased cytosolic pH, K+, and anions. Journal of Experimental Botany 66, 813–825.

Rodriguez PL, Lozano-Juste J, Albert A. 2019. PYR/PYL/RCAR ABA receptors. In: Seo M, Marion-Poll A, eds. Advances in Botanical Research, Academic Press 92, 51–82.

Ruiz-Partida R, Prado S, Villarroya M, Velázquez-Campoy A, Bravo J, Armengod ME. 2018. An Alternative Homodimerization Interface of MnmG Reveals a Conformational Dynamics that Is Essential for Its tRNA Modification Function. Journal of Molecular Biology 430, 2822–2842.

Saez A, Rodrigues A, Santiago J, Rubio S, Rodriguez PL. 2008. HAB1-SWI3B interaction reveals a link between abscisic acid signaling and putative SWI/SNF chromatin-remodeling complexes in Arabidopsis. Plant Cell 20, 2972–2988.

Safronov O, Kreuzwieser J, Haberer G, et al. 2017. Detecting early signs of heat and drought stress in *Phoenix dactylifera* (date palm). PLoS ONE 12, e0177883.

Santiago J, Rodrigues A, Saez A, Rubio S, Antoni R, Dupeux F, Park SY, Márquez JA, Cutler SR, Rodriguez PL. 2009. Modulation of drought resistance by the abscisic acid receptor PYL5 through inhibition of clade A PP2Cs. Plant Journal 60, 575–588.

Schweighofer A, Hirt H, Meskiene I. 2004. Plant PP2C phosphatases: Emerging functions in stress signaling. Trends in Plant Science 9, 236–243.

Seiler C, Harshavardhan VT, Reddy PS, et al. 2014. Abscisic acid flux alterations result in differential abscisic acid signaling responses and impact assimilation efficiency in barley under terminal drought stress. Plant Physiology 164, 1677–1696.

Sharp RE, Poroyko V, Hejlek LG, Spollen WG, Springer GK, Bohnert HJ, Nguyen HT. 2004. Root growth maintenance during water deficits: Physiology to functional genomics. Journal of Experimental Botany 55(407), 2343–2351.

Torres MF, Mathew LS, Ahmed I, Al-Azwani IK, Krueger R, Rivera-Nuñez D, Mohamoud YA, Clark AG, Suhre K, Malek JA. 2018. Genus-wide sequencing supports a two-locus model for sex-determination in *Phoenix*. Nature Communications 9, 3969.

Umezawa T, Sugiyama N, Mizoguchi M, Hayashi S, Myouga F, Yamaguchi-Shinozaki K, Ishihama Y, Hirayama T, Shinozaki K. 2009. Type 2C protein phosphatases directly regulate abscisic acid-activated protein kinases in Arabidopsis. Proceedings of the National Academy of Sciences of the United States of America 106, 17588–17593.

Vaidya AS, Helander JDM, Peterson FC, et al. 2019. Dynamic control of plant water use using designed ABA receptor agonists. Science 366.

Vaidya AS, Peterson FC, Yarmolinsky D, et al. 2017. A Rationally Designed Agonist Defines Subfamily IIIA Abscisic Acid Receptors As Critical Targets for Manipulating Transpiration. ACS Chemical Biology 12, 2842–2848.

Vlad F, Rubio S, Rodrigues A, Sirichandra C, Belin C, Robert N, Leung J, Rodriguez PL, Laurière C, Merlot S. 2009. Protein phosphatases 2C regulate the activation of the Snf1-related kinase OST1 by abscisic acid in Arabidopsis. Plant Cell 21, 3170–3184.

Wang X, Guo C, Peng J, et al. 2019. ABRE-BINDING FACTORS play a role in the feedback regulation of ABA signaling by mediating rapid ABA induction of ABA co-receptor genes. New Phytologist 221, 341–355.

Witte CP, Noël LD, Gielbert J, Parker JE, Romeis T. 2004. Rapid one-step protein purification from plant material using the eight-amino acid StrepII epitope. Plant Molecular Biology 55, 135–147.

Xiao TT, Raygoza AA, Pérez JC, et al. 2019. Emergent protective organogenesis in date palms: A morpho-devo-dynamic adaptive strategy during early development. Plant Cell 31, 1751–1766.

Yaish MW, Patankar H. V., Assaha DVM, Zheng Y, Al-Yahyai R, Sunkar R. 2017. Genomewide expression profiling in leaves and roots of date palm (*Phoenix dactylifera* L.) exposed to salinity. BMC Genomics 18, 246.

Yang Z, Liu J, Tischer S V., Christmann A, Windisch W, Schnyder H, Grill E. 2016. Leveraging abscisic acid receptors for efficient water use in Arabidopsis. Proceedings of the National Academy of Sciences of the United States of America 113(24), 6791–6796.

Yang Z, Liu J, Poree F, et al. 2019. Abscisic Acid Receptors and Coreceptors Modulate Plant Water Use Efficiency and Water Productivity. Plant Physiology 180, 1066–1080.

Yin Y, Zhang X, Fang Y, et al. 2012. High-throughput sequencing-based gene profiling on multi-staged fruit development of date palm (*Phoenix dactylifera*, L.). Plant Molecular Biology 78, 617–626.

Zhao Y, Chan Z, Gao J, et al. 2016. ABA receptor PYL9 promotes drought resistance and leaf senescence. Proceedings of the National Academy of Sciences of the United States of America 113, 1949–1954.

